# Antibiofilm activity of Fmoc-phenylalanine against Gram-positive and Gram-negative bacterial biofilms

**DOI:** 10.1101/2020.08.03.232629

**Authors:** Himanshi Singh, Avinash Y. Gahane, Virender Singh, Shreya Ghosh, Ashwani Kumar Thakur

## Abstract

**Background:** Biofilm associated infections are the major contributor of mortality, morbidity and financial burden in patients with bacterial infection. Molecules with surfactant behaviour are known to show significant antibiofilm effect against these infections. Thus, newly discovered antibacterial Fmoc-phenylalanine (Fmoc-F) and other Fmoc-amino acids (Fmoc-AA) with surfactant properties, could have potential antibiofilm properties.

**Objectives:** To evaluate and characterise the antibiofilm activity of Fmoc-F and some Fmoc-AA against various clinically relevant bacteria.

**Methods:** Biofilm inhibition and eradication was evaluated by crystal violet staining procedure along with scanning electron microscopy (SEM). Attenuated Total Reflection - Fourier Transform Infrared Spectroscopy (ATR-FTIR), Biochemical assays and Congo red staining were employed to investigate mechanism of antibiofilm action.

**Results:** We showed that Fmoc-F not only inhibits the biofilm formation in *S. aureus* and *P. aeruginosa*, but also eradicates the already formed biofilms over the surface. Further, Fmoc-F coated glass surface resists *S. aureus* and *P. aeruginosa* biofilm formation and attachment, when biofilm is grown over the surface. The mechanistic investigation suggests that Fmoc-F reduces the ECM components such as proteins carbohydrates and eDNA in the biofilm and affect its stability *via* direct interactions with ECM components and/ or indirectly through reducing bacterial cell population. Finally, we showed that Fmoc-F treatment in combination with other antibiotics such as vancomycin and ampicillin synergistically inhibit biofilm formation.

**Conclusions:** Overall, the study demonstrates the potential application of Fmoc-F and other Fmoc-AA molecules individually as well as in combination as antibiofilm agents and antibiofilm coating material for treating biofilm associated infections.

## Introduction

Bacterial Biofilms are the complex bacterial aggregates usually encased within the secreted extracellular matrix (ECM).^1^ Several pathogenic and non-pathogenic bacteria form biofilms in response to multiple environmental stresses that involve altered regulatory pathways and gene expression changes. The biofilm cells adhere to the different biotic and abiotic surfaces, in contrast to planktonic cells which are generally free flowing bacteria. The ECM which is composed of polysaccharides, proteins and nucleic acids provides stability to the biofilm cells and mediates surface adhesion.^2^ These bacterial biofilms are mainly associated with chronic and persistent infections, as well as nosocomial infections such as medical device or implants related infections. According to National Institute of Health (NIH), 65% of microbial and 85% of all chronic infections are caused due to biofilm formation.^3^ These include bacterial vaginosis, dental plaques, gingivitis, meningitis, endocarditis, rhinosinusitis, osteomyelitis, non-healing chronic wounds, urinary tract infections, middle ear infections and prosthesis & implantable device- related infections in humans.^4^ Biofilm infections are extremely difficult to treat due to lack of diagnostic methods, biomarkers and specific antibiotics.^5^ Most of the antibacterial drugs become less effective when they are used against biofilm associated bacteria. Moreover, with the disparity of antibiotic concentration throughout the biofilm, bacterial cells are sometimes exposed to concentration below inhibitory level due to which they develop antibiotic resistance.^6,7^

Different strategies have been utilised either to combat biofilms formation or eradicate already formed bacterial biofilms. While antimicrobial drugs such as Thioridazine and Phenylalanine-arginine-naphthylamide directly affect biofilm cells,^8–10^ antimicrobial peptides or matrix disrupting enzymes such as DNase and Amylase can be utilised to enhance ECM penetration of other antibiotics and show synergistic effect to eradicate bacterial biofilms.^7,10,11^ Recently, nanoparticles which develop shear stress in biofilms and enhanced membrane permeability shown to be effective against *S. aureus* and *P. aeruginosa* biofilms.^12^ In spite of good efficiency of various antimicrobial peptides, enzymes, nanoparticles etc. towards different biofilm forming bacteria, each having their own limitations such as low metabolic stability, toxicity, high cost and poor scalability. Hence, small molecules are being examined for their potential applications as antimicrobials and biofilm disruptors. For example, D-amino-acids and their mixtures shown to inhibit biofilm formation in *B. subtilis*, *S. aureus* and *P. aeruginosa* infections.^13^ It was concluded that D-amino acids had effect on cell adhesion proteins which prevents assembly of cells on the surface.^14^ Recently, McCloskey *et al.* showed that fluorenylmethoxycarbonyl (Fmoc) conjugated peptides (Fmoc-peptide) such as Fmoc-FF (di-phenylalanine), Fmoc-FFKK (di-lysine) and Fmoc-FFOO (di-ornithine) can eradicate both Gram-positive (*S. aureus, S. epidermis*) and Gram-negative (*P. aeruginosa, E. coli*) bacteria biofilm.^15^ These Fmoc-peptides possess optimal hydrophobicity (40-60%) and surfactant like properties to cause sufficient interaction with bacterial membrane for cell lysis.

Recently, we have shown that Fmoc-phenylalanine (Fmoc-F) and other Fmoc-amino acids possess good antibacterial activity which is associated with their surfactant like properties.^16^ Further, molecule with surfactant properties have been known to eradicate biofilm and enhance antimicrobial efficacy in biofilm.^17–19^ Thus, present study is undertaken to investigate the effect of Fmoc-F on biofilm formation by clinically relevant Gram-positive and Gram-negative bacteria.

## Materials and methods

### Bacterial strains and culture conditions

Different bacterial strains such as *S. aureus* (MTCC 737) and *P. aeruginosa* (MTCC 4673) were purchased from Institute of Microbial Technology (IMTECH, Chandigarh, India). For all the experiments, primary culture was prepared by growing a single colony in Tryptone Soya Broth (TSB) media and used to obtain secondary culture (OD = 0.3 to 0.6). Fmoc-F and other Fmoc-AA solutions were prepared as described by Gahane *et al.*, 2018. Briefly, Fmoc-F was dispersed in 50 mM phosphate buffer (PB) to obtain required concentration. The suspension was sonicated for 30 min and then heated at 60 °C to obtain a uniform solution. Further details about experimental procedure are given in supporting information.

## Results and discussion

### Fmoc-F inhibits biofilm formation in Gram-positive as well as Gram-negative bacteria

Several Gram-positive and Gram-negative bacterial infections involve biofilm formation which makes the infection chronic and antibiotic resistant. Thus, initially we checked effect of Fmoc-F treatment on biofilm formation by Gram-positive bacteria such as *S. aureus*. Incubation of *S. aureus* with Fmoc-F (1.2 mM; Minimum Bactericidal Concentration) for 48 h significantly reduced the formation of bacterial biofilm, as shown by decrease in crystal violet absorbance compare to the control (*p* ≤ 0.05, Figure 1a). We expected the less biomass formation, as Fmoc-F with 1.2 mM concentration is able to kill *S. aureus* and reduces cell number in media.^16^ To increase the spectrum of Fmoc-F activity, we checked its effect against biofilm formation by Gram-negative bacteria i.e. *P. aeruginosa*. Incubation of *P. aeruginosa* with Fmoc-F (1.2 mM) for 48 h also resulted in significant reduction in bacterial biofilm formation (*p* ≤ 0.05, Figure 1a). Bacterial culture consists of planktonic cells suspended in the media and biofilm cells encased in dense ECM attached to the surface. These cell types exhibit varying susceptibility to a particular antibiotic due to difference in physiology, population density and surrounding environment.^20^

**Figure 1.**
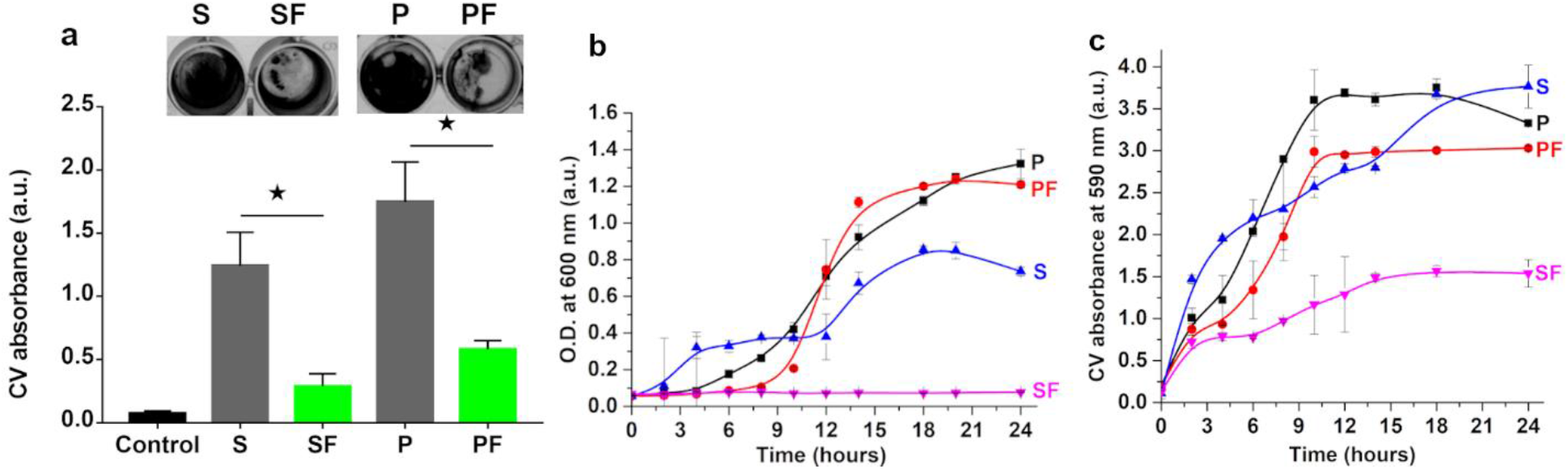
Fmoc-F treatment reduces the bacterial biofilm formation. a) Inhibition of biofilm formation of *S. aureus* and *P. aeruginosa* by Fmoc-F (1.2 mM) treatment shown by crystal violet staining assay. b) Growth profile of planktonic cells of *S. aureus* and *P. aeruginosa* in media in the presence of Fmoc-F (1.2 mM). c) Growth profile of biofilm cells and associated biomass of *S. aureus* and *P. aeruginosa* biofilms grown in presence of Fmoc-F (1.2 mM). \**p* ≤ 0.05 *vs* control, student *t*-test. S - *S. aureus* biofilm, P - *P. aeruginosa* biofilm, SF - *S. aureus* biofilm in presence of Fmoc-F, PF - *P. aeruginosa* biofilm in presence of Fmoc-F.

Thus, to investigate whether the observed reduction in biofilm formation is *via* planktonic cell reduction or due to direct effect of Fmoc-F on biofilm cells and associated ECM matrix (Biomass), we assessed the growth profiles of planktonic cells and biomass in *S. aureus* and *P. aeruginosa*. In control groups, a small lag phase was observed in planktonic cells growth for both bacteria, followed by logarithmic growth phase after 2-4 h (Figure 1b). However, treatment with Fmoc-F (1.2 mM) resulted in no increase in planktonic cell number of *S. aureus* at all, while *P. aeruginosa* planktonic cells showed initial lag phase of 8 h followed by logarithmic growth phase similar to that of control. In case of biofilm cells, the control group showed steady increase in the cell attachment throughout the logarithmic phase and after 12 h, a constant amount of biomass was attached. However, in presence of Fmoc-F, the biomass formation was less and not robust as compared to the control in both bacteria and could not sustain washing due to decreased adhesiveness (Figure 1c). The effect was robust in case of *S. aureus*. Interestingly, Fmoc-F did not affect planktonic cell number of *P. aeruginosa*, but the final biomass attachment on the surface was found to be less. These results suggest that Fmoc-F reduces biofilm formation in Gram-positive bacteria synergistically, by inhibiting planktonic as well as biofilm cell growth and their interaction with ECM components. However, in case of Gram-negative bacteria, Fmoc-F might affect ECM stability *via* direct interactions with matrix components such as adhesion proteins making the biofilm loosely held on the surface which leads to biofilm disintegration and reduced biofilm formation.

### Fmoc-F promotes biofilm eradication in Gram-positive and Gram-negative bacteria

In clinical settings, medical device-related biofilm infections often involve the presence of already formed biofilms. Examples include various implants such as cardiac pacemakers and prosthetic heart valves, orthopaedic devices and catheters. These biofilm infections are difficult to treat and in most cases the device is removed and replaced, which is a very risky and costly procedure. Thus, recent scientific efforts are focussed on eradication of already formed biofilm on the surface along with prevention of its formation. Synthetic and natural surfactants have been implicated for inhibition and eradication of preformed biofilms.^18, 21^ They lower the surface tension and adhesiveness of biofilm with surface, loosen the matrix and disperse it. Previously, we showed that Fmoc-F also possess mild surfactant properties and ability to reduce surface tension. Thus, we tested Fmoc-F for its ability to eradicate already existing (preformed) biofilm. Preformed biofilms of *S. aureus* and *P. aeruginosa* were grown after 48 h of incubation period in 96 wells plates and treated with Fmoc-F (1-3 mM) for 2 h. The attached biomass to the polystyrene surface was quantified with crystal violet staining. Results showed that treatment with ≥ 1.2 mM of Fmoc-F significantly eradicates the adhered biofilm from the surface in both bacteria (Figure 2a and 2b). Interestingly, the minimum concentration for biofilm eradication is nearly equal to the critical micelle concentration (CMC) of Fmoc-F (1.25 mM), which suggests that Fmoc-F eradicates the biofilm through micelle formation.^16^ Since the anti-biofilm effect was observed in both bacteria, Fmoc-F activity is independent of cell type and could be mediated through its interaction with biofilm ECM. We speculated that, beyond CMC, Fmoc-F forms micelles which trap hydrophobic ECM components and cell debris just like surfactants that help in removing attached biomass. Also, micelles provide external shear forces to the overall matrix leading to dispersion of biofilm components.^22^

**Figure 2.**
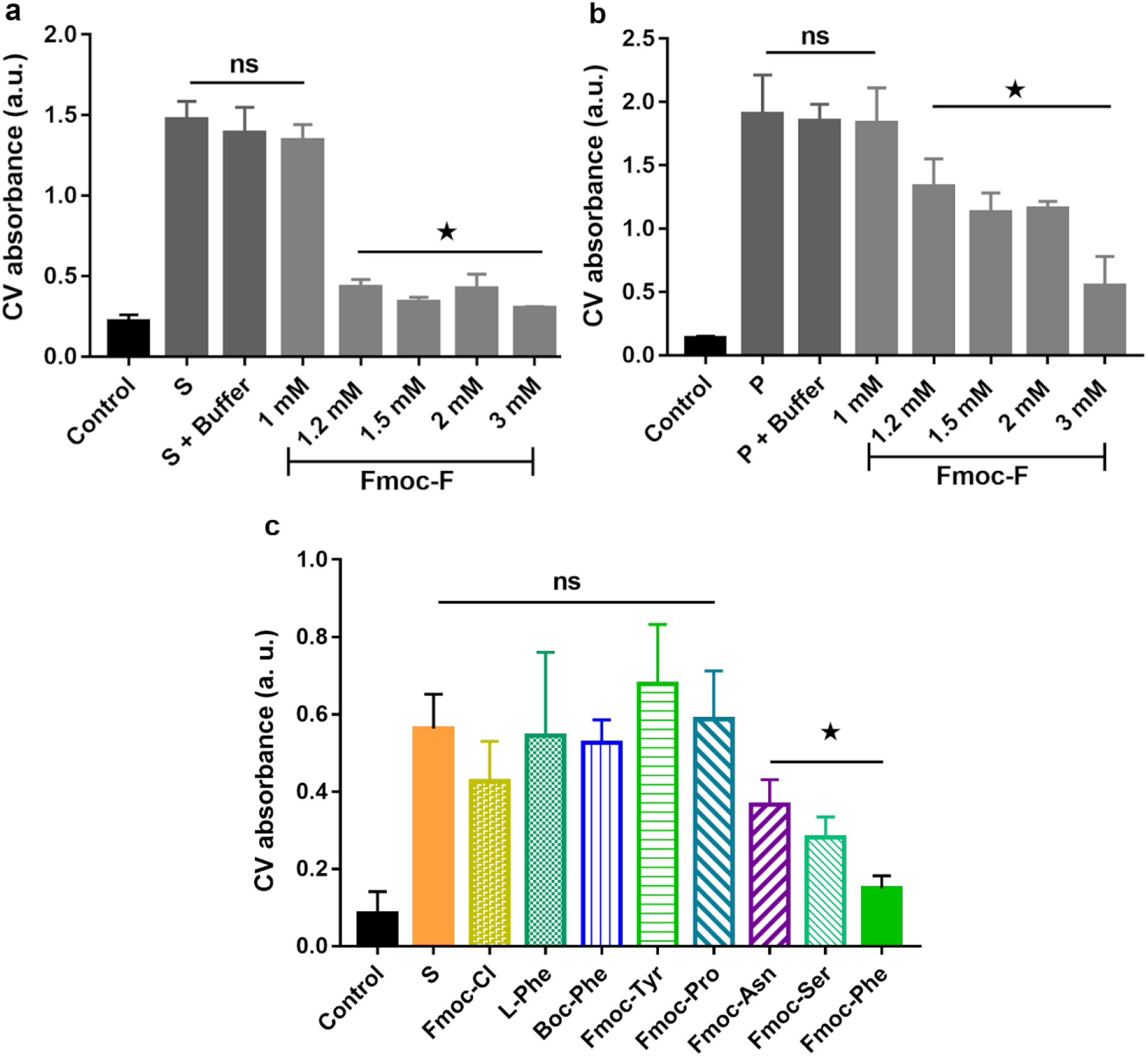
Fmoc-F treatment eradicates preformed bacterial biofilms as shown by crystal violet staining assay a) Eradication of *S. aureus* biofilm by different concentrations of Fmoc-F (1 - 3 mM). b) Eradication of *P. aeruginosa* biofilm by different concentrations of Fmoc-F (1 - 3 mM). c) Eradication of *S. aureus* biofilm by L-phenylalanine derivatives and Fmoc-amino acids. \**p* ≤ 0.05 *vs* control, One-way ANOVA followed by Dunnett’s multiple comparison test. ns – Non-significant

To understand the structure–activity relationship, we tested various Fmoc-F derivatives (1.5 mM) against *S. aureus* biofilms. Interestingly, individual treatment with Fmoc-Cl or L-Phe resulted in complete loss of anti-biofilm activity (Figure 2c, Figure S1). This suggests that covalent linkage between Fmoc moiety and L-Phe is essential for the activity. Loss of activity due to replacement of Fmoc with N-(tert-Butoxycarbonyl) (Boc) group suggests, presence of more bulky groups with added hydrophobicity is also essential for anti-biofilm activity. Further, to understand the significance of amino acid substitution, we tested other Fmoc-AA for their anti-biofilm activity. We found that Fmoc-Ser and Fmoc-Asn exhibited biofilm eradication activity similar to that of Fmoc-F, while loss of activity was observed for Fmoc-Pro and Fmoc-Tyr (Figure 2c, Figure S1).

Interestingly, Fmoc-AA exhibit a similar pattern as their antibacterial activity, except Fmoc-Asn.^16^ The inactivity of Fmoc-Pro can be attributed to its rigid, weakly hydrophobic and neutral side chain which might affect the intensity of shear force acting on the cells and matrix.^23^ While absence of anti-biofilm activity in Fmoc-Tyr suggest that apart from aromaticity, overall hydrophobicity of molecule (bulky functional group and attached amino acids) is important for the activity. That is why some effect (insignificant) is seen in Fmoc-Cl (Figure 2c, Figure S1). Overall results suggest that enhanced hydrophobicity of Fmoc-AA with specific structural gain is essential for biofilm interaction and its disintegration.

### Fmoc-F coated surfaces inhibit biofilm formation and attachment

Prevention of biofilm formation is considered to be easier, safer and more cost-effective way than treating a biofilm that has been already formed. Several researchers utilised modified surfaces that are resistant to the biofilm formation or used coating surfaces with anti-biofilm agents to prevent biofilm formation. Fmoc-F has the ability to self-assemble itself and form a transparent hydrogel,^24^ which could be utilised for anti-biofilm coating. To test this, Fmoc-F coated coverslips were prepared by overnight incubation with Fmoc-F solution (15 mM) and tested for biofilm formation. The Fmoc-F coated coverslip were translucent in appearance (Figure 3a) and showed presence of small Fmoc-F self-assemblies on the surfaces (Figure 3b). Also, Incubation of *S. aureus* and *P. aeruginosa* over the Fmoc-F coated coverslip resulted in significant decrease in bacterial biofilm formation as compare to control coverslip without any coat (Figure 3c). This was further confirmed by scanning electron microscopic (SEM) image of Fmoc-F coated coverslip used to grow *P. aeruginosa* biofilm (Figure 3d). These results suggest that Fmoc-F could be used as a coating material for various devices and equipments to prevent the bacterial biofilm formation on the various surfaces. As hydrophilic environment on the surface is required for biofilm attachment,^25^ it might be possible that coating with hydrophobic Fmoc-F reduces the surface wettability of coverslip and hence biomass attachment. Also, the presence of Fmoc-F on the surface might interfere with the initial interactions necessary for formation of ECM conditioning film required for biofilm surface adhesion and stability.^26^

**Figure 3.**
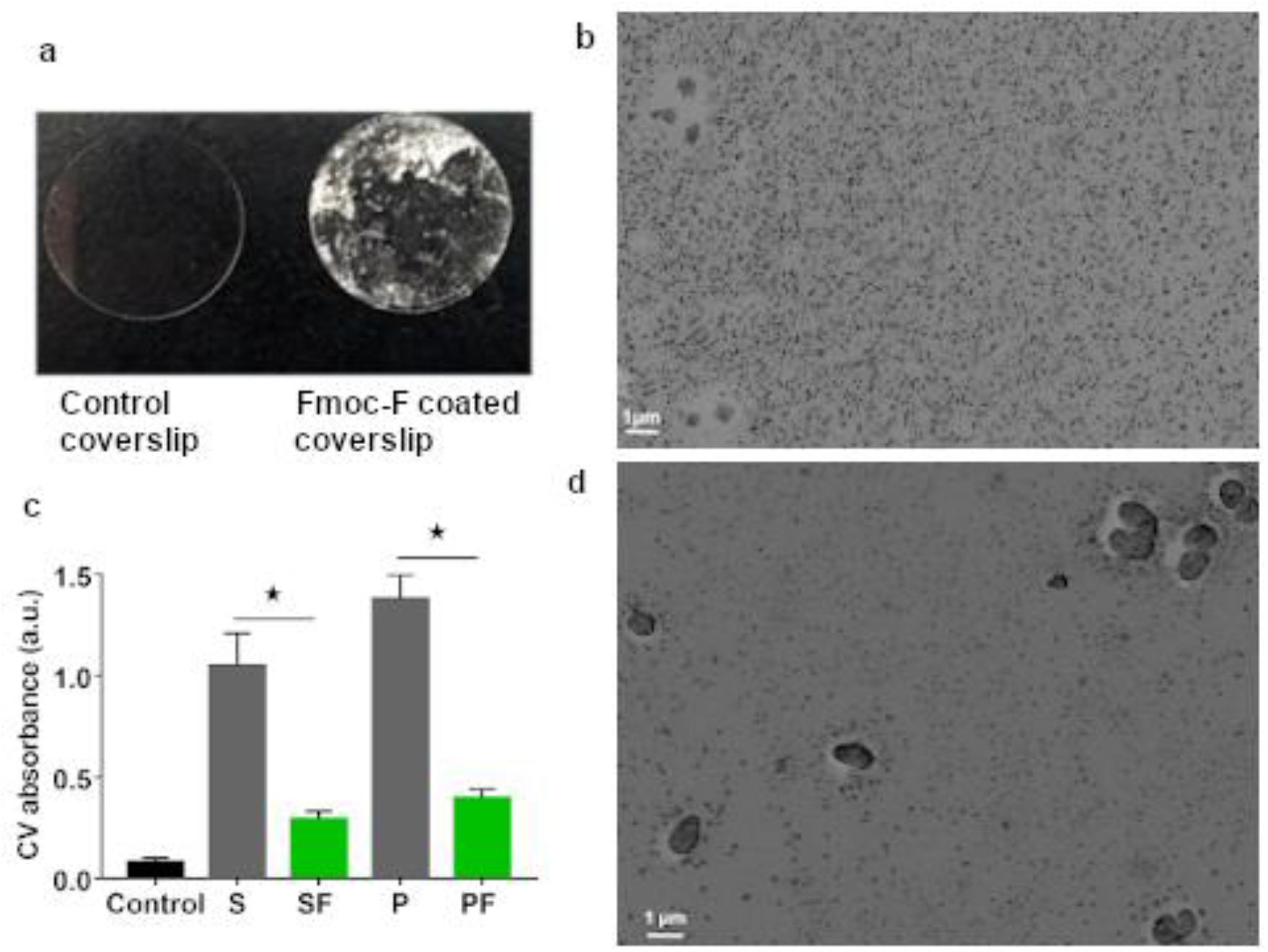
Fmoc-F coated surfaces inhibit biofilm formation and attachment. a) Digital images of glass coverslip coated with Fmoc-F solution (15 mM). b) SEM image of glass coverslip coated with Fmoc-F solution (15 mM). c) Inhibition of biofilm formation of *P. aeruginosa* by Fmoc-F coated surface of glass coverslip. d) SEM image of *P. aeruginosa* cells grown on Fmoc-F coated glass coverslip. \**p* ≤ 0.05 *vs* control, student *t*-test. Magnification - 5000 X. S - *S. aureus* biofilm on uncoated coverslip, P - *P. aeruginosa* biofilm on uncoated coverslip, SF - *S. aureus* biofilm on Fmoc-F coated coverslip, PF - *P. aeruginosa* biofilm on Fmoc-F coated coverslip

### Fmoc-F treatment reduces various ECM component in the bacterial biofilms

The integrity of bacterial biofilm is mainly dependent on ECM properties which makes the biofilm cells connected and protected from the surrounding environment. The major ECM components secreted by the bacterial cells include proteins, carbohydrates, lipids and eDNA where they perform collective functions for growth and stability of biofilms.^27^ Since, Fmoc-F inhibits the biofilm formation and eradication by Gram-negative bacteria with minimum effect on planktonic cell population (Figure 1b), we thought that Fmoc-F action is not cell specific and might be mediated through its interaction with various ECM components. Thus, to check this, we studied differences in ECM components from biofilms formed in the presence or absence of Fmoc-F using ATR-FTIR analysis. For quantitative analysis, sodium-azide was used as an internal standard. The primary FTIR spectra of ECM obtained from biofilm of untreated bacteria (control) showed major peaks at ~3400-2900 cm^−1^ representing lipids, ~1800-1500 cm^−1^ representing proteins, ~1400-1200 cm^−1^ representing nucleic acids & phosphorylated proteins and ~1100-900 cm^−1^ representing polysaccharides in the ECM (Figure 4).^28^ FTIR spectra of ECM derived from biofilms of *S. aureus* and *P. aeruginosa* grown in the presence of Fmoc-F (1 mM or 1.2 mM) also showed similar peaks to that of control, suggesting presence of similar ECM components. However, there was approximately 40% decrease in proteins, 50% decrease in polysaccharides and around 60% decrease in nucleic acid components compared to control ECM as suggested by decrease in intensity of respective spectral peaks (Figure 4a and 4b).

**Figure 4.**
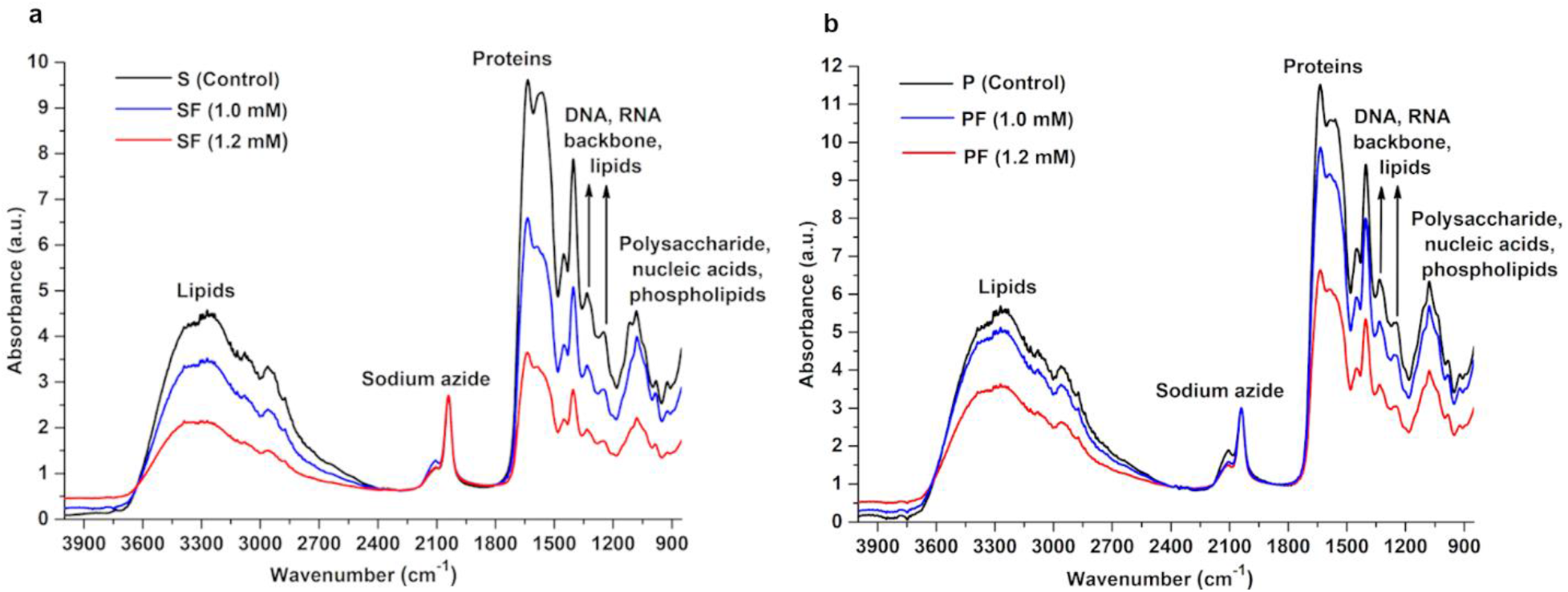
ATR-FTIR spectra of ECM extracted from bacterial biofilms. a) FTIR spectra of ECM extracted from *S. aureus* biofilm formed in the presence of Fmoc-F (1 and 1.2 mM). b) FTIR spectra of ECM extracted from *P. aeruginosa* biofilm formed in the presence of Fmoc-F (1 and 1.2 mM).

To confirm these results, we quantified the ECM components in the biofilm formed in presence of varying Fmoc-F concentrations (0.5 – 1.2 mM) using different biochemical assays. Results showed that, Fmoc-F dose dependently reduced the protein content in the ECM compared to the control i.e. without Fmoc-F treatment (Figure 5a, Figure S2). The observed effect was significant in *S. aureus* and *P. aeruginosa* biofilms when incubated with 1 mM and 0.5 mM of Fmoc-F, respectively. The decreased protein content might result in unstable biofilm due to reduced ionic and surface interactions. Also, ECM proteins ensure proper cell adhesion and quorum sensing among bacteria for biofilm formation,^29–31^ which might be compromised in presence of Fmoc-F. Similarly, we observed significant reduction in polysaccharides as well as nucleic acid content in the ECM of the biofilms grown in the presence of Fmoc-F (Figure 5b and 5c). Fmoc-F treatment (1.2 mM) significantly reduced the carbohydrate content in ECM from *S. aureus* and *P. aeruginosa* biofilms and DNA content from *S. aureus* biofilms. DNA and polysaccharides are involved in biofilm structuring and resistance from external forces, respectively.^29^ Overall results confirm that, Fmoc-F treatment reduces ECM components such as proteins, polysaccharides and nucleic acids in Gram-positive and Gram-negative bacterial biofilms and might be the reason behind formation of mechanically unstable and compromised biofilms.

**Figure 5.**
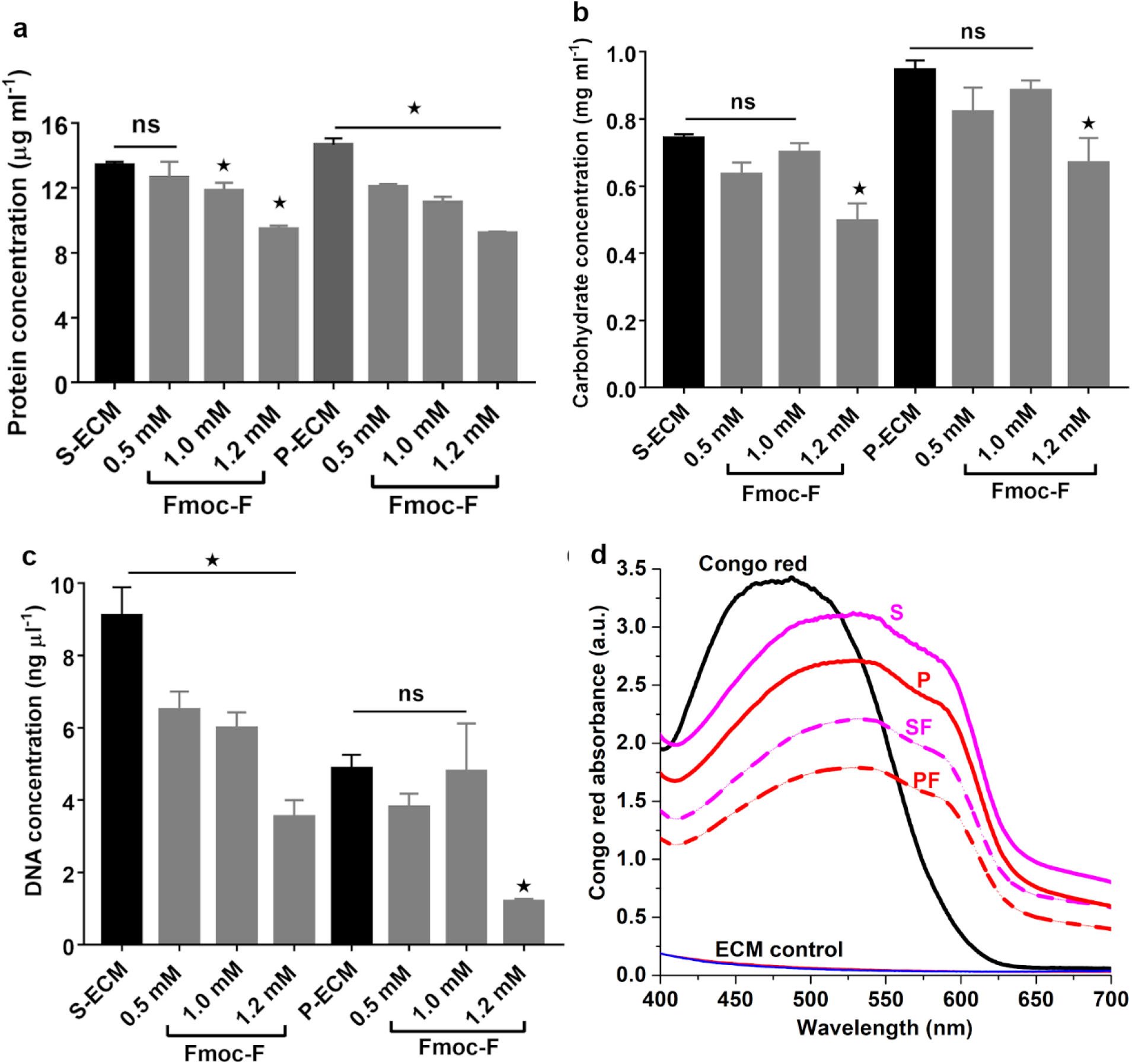
Effect of Fmoc-F (0.5, 1 and 1.2 mM) on various ECM components a) proteins b) carbohydrates c) DNA. \**p* ≤ 0.05 *vs* control, student *t*-test, ns – Non-significant. d) Effect of Fmoc-F on amyloid structures in ECM: Graph showing Congo red absorbance spectra in the presence of ECM extracted from *P. aeruginosa* biofilm formed with Fmoc-F (1.2 mM). S - *S. aureus* ECM, P - *P. aeruginosa* ECM, SF - *S. aureus* ECM treated with Fmoc-F, PF - *P. aeruginosa* ECM treated with Fmoc-F.

### Effect of Fmoc-F on amyloid structures

In order to form a mature biofilm, structuring of various ECM components is essential for mechanical stability. During biofilm maturation, amyloids formation has recently emerged out as an important event. Basically, different adhesion proteins as well as other ECM proteins form complex amyloid-like structures which reduce antibiotic penetrance and make biofilms resistant to the antibiotics.^32–35^ In this context, effect of Fmoc-F on amyloid structures in ECM was studied using Congo-red absorption assay. In Figure 5d, spectral shift of Congo red peak (from 480 nm to 540 nm) in presence of ECM indicates presence of amyloid-like structures in *S. aureus* and *P. aeruginosa* biofilms.^36^ The reduced absorbance intensity of Congo red in the presence of Fmoc-F (1.2 mM) treated ECM from both bacteria, suggest that Fmoc-F reduces the amyloid formation or impart structural changes in the amyloids (Figure 5d). The observed effect was found to be in direct correlation with Fmoc-F incubation time with ECM (Figure S3). This was further confirmed by ThT binding assay, where Fmoc-F treated ECM showed reduced ThT fluorescence intensity compared to untreated ECM (Figure S3). These results suggest that Fmoc-F is able to reduce amyloid formation in the ECM of bacterial biofilms which might be responsible for reduced biofilm integrity and mechanical stability.^37,38^

### Combination treatment of Fmoc-F with different antibiotics produce synergistic effect

In clinical settings, biofilm infections are usually treated with combinations of various antibiotics.^39^ It is also an alternative approach to lower the required dose of antibiotics and obtain the possible synergistic effect.^40^ Thus, we checked the biofilm eradication effect of Fmoc-F given in combination with vancomycin or ampicillin. Vancomycin and ampicillin are the most common clinical and veterinary antibiotics used to treat bacterial infections. However, their efficacy to treat biofilm associated infections is 50 - 200 times lower than that of other infections.^41^ Firstly, we incubated *S. aureus* or *P. aeruginosa* biofilms with different concentrations of either vancomycin (1 - 35 μM), ampicillin (4.4 – 143 μM) or Fmoc-F (37 - 1200 μM) and found that vancomycin ≤ 2 μM or ampicillin ≤ 36 μM or Fmoc-F ≤ 600 μM non-significantly eradicate the preformed bacterial biofilms (Figure S4). Next, to examine the effect of Fmoc-F combinatorial treatment on biofilm eradication, we incubated three sub-effective doses of Fmoc-F (150, 75, 37.5 μM) with sub-effective dose of vancomycin (1.09 μM) or ampicillin (4.45 μM). All the tested combinations showed better biofilm eradication effect than individual treatment in both the bacteria, as shown by decrease in crystal violet absorption (Figure 6). There was 32-fold and 8-fold reduction in effective dose for Fmoc-F (1.2 mM to 37.5 μM) and ampicillin (36.25 μM to 4.45 μM), respectively (Figure 6 and S4). Generally, fourfold reduction of the effective dose of the antibiotic is considered as a synergistic effect.^42^ These results suggest that Fmoc-F could be used in combination with other antibiotics against biofilm associated infections to produce synergistic effect. We think that by affecting ECM integrity, Fmoc-F might improve the penetration of other antibiotics given in combination and exhibit synergy. The complementary action of Fmoc-F with antibiotics can be an effective solution against the growing threat of antibiotic resistance of bacterial biofilms.

**Figure 6.**
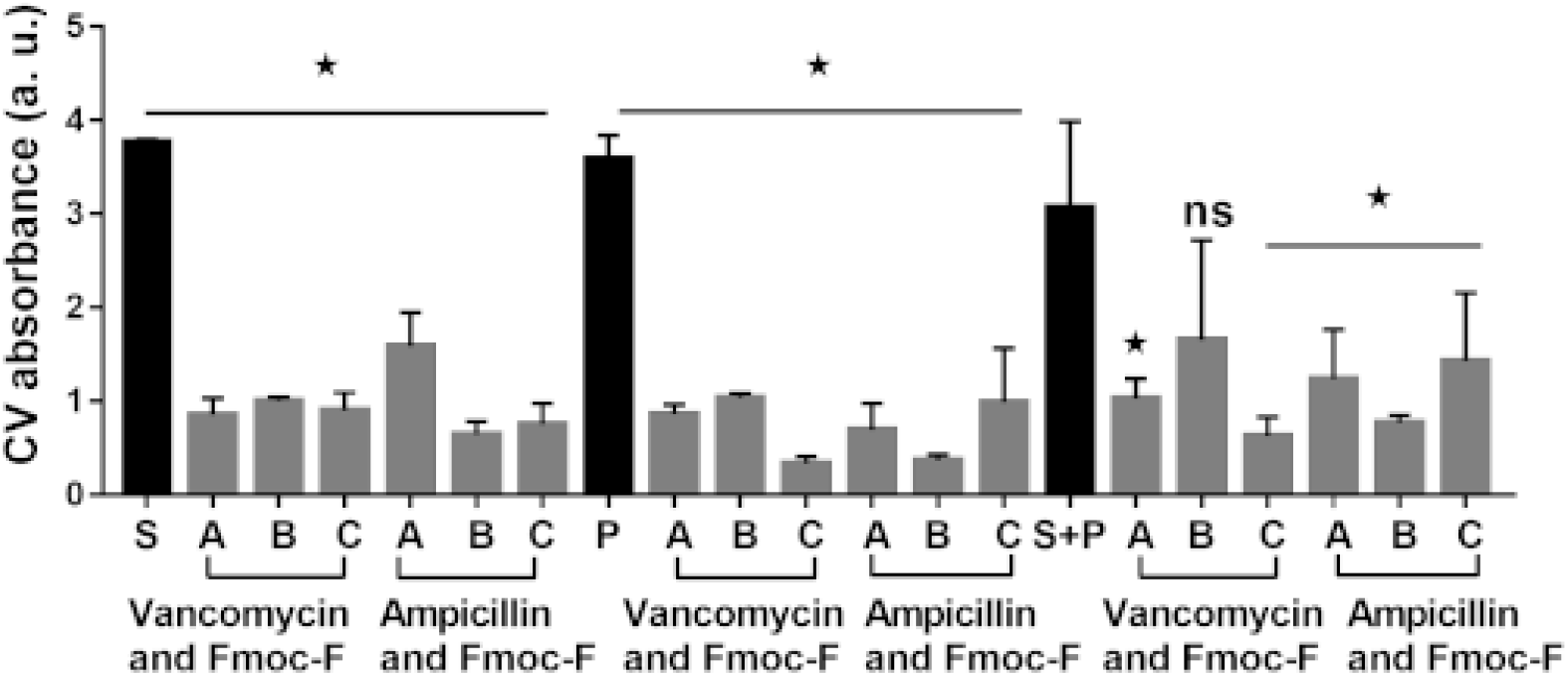
Eradication of *S. aureus, P. aeruginosa* and co-culture biofilm by combination treatment of Fmoc-F (150, 75, 37.5 μM) with vancomycin (1.09 μM) or ampicillin (4.45 μM). Vanco - Vancomycin, Amp - Ampicillin, A,B,C - different concentrations of Fmoc-F, S+P - co-cultured biofilm of both *S. aureus* and *P. aeruginosa*. \**p* ≤ 0.05 *vs* control, One-way ANOVA followed by Dunnett’s multiple comparison test. ns – Non-significant

## Conclusion

The present study characterises the anti-biofilm activity of Fmoc-F molecule against clinically relevant biofilm forming Gram-positive and Gram-negative bacteria. Fmoc-F is found to be effective in preventing biofilm formation as well as eradicate mature biofilms from attached surfaces. In case of Gram-positive bacteria such as *S. aureus*, Fmoc-F prevents biofilm formation by reducing cell population *via* its antibacterial effect. Interestingly, Fmoc-F also inhibits biofilm formation by Gram-negative bacteria such as *P. aeruginosa* which suggests that Fmoc-F affects ECM matrix composition and stability through direct interactions with ECM components. Eradication of already formed biofilms and reduction in ECM components like eDNA, polysaccharides and proteins in Gram-positive as well as Gram-negative bacterial biofilms by Fmoc-F treatment further supports this hypothesis. We think that Fmoc-F directly interacts with ECM components and alters its composition thereby inhibits the structuring and crosslinking of ECM resulting in mechanically unstable biofilm. Considering the surfactant like behaviour, Fmoc-F molecules could exert shear stress and disrupt mature biofilms via formation of micelle like structures. To extend the applicability, we further showed that Fmoc-F coated surfaces and its combination with other antibiotics synergistically inhibit biofilm formation. Overall, this work establishes the broad-spectrum, biofilm inhibition activity of Fmoc-F and its biomedical application as anti-biofilm coating material. It is known that different quorum sensing pathways and operons play essential role in establishing stable biofilm through cell communication. As Fmoc-F targets ECM of biofilm, molecular mechanism of action can be explored in future.

## Acknowledgement

Authors thank IITK, CSIR and MHRD for funding the fellowships.

## Transparency declarations

None to declare.

