## Supplementary Information for "Antibiofilm activity of Fmoc-phenylalanine against Gram-positive and Gram-negative bacterial biofilms"

**Contents**

1. Materials and methods
2. Anti-biofilm activity of Fmoc- amino acids and L-phenylalanine derivatives against *S. aureus* biofilm S1
3. Standard curves for biochemical assays S2
4. Effect of Fmoc-F on amyloid structures in ECM S3
5. Eradication of *S. aureus* and *P. aeruginosa* biofilm by treatment of different concentrations of antibiotics S4
6. References

### Materials and methods

#### Crystal violet staining

Biofilm mass attached on different surfaces were quantified by crystal violet staining as described by O'Toole in 2011. Briefly, 0.1% of crystal violet solution was added to the different biofilm samples. After 30 min, the solution was decanted and biofilms were washed twice using water and air dried. Crystal violet attached to biofilm was dissolved in 30% acetic acid for 30 minutes and absorbance was recorded at 590 nm using multimode plate reader (EnSpire).[1]

#### Biofilm susceptibility assay

To observe the effect of Fmoc-F against biofilm formation, Fmoc-F (1.2 mM, 20 mL) in TSBg (TSB supplemented with 0.01% glucose) sterile media was prepared by adding sufficient quantity of previously prepared Fmoc-F solution in PB. To this, 20 µL of secondary culture (OD ~ 0.6) from *S. aureus* or *P. aeruginosa* was added. Subsequently, 200 µL of culture in Fmoc-F containing media was transferred to 96-well microtiter plate. The plate was properly covered with its lid and incubated at 37$^{o}$C in static condition. After 24 hours, media was replaced with fresh media with Fmoc-F. After 48 hours incubation period, unattached planktonic cells were decanted by inverting the plate and washed vigorously twice using 1X PBS. The attached biomass was quantified by crystal violet staining as mentioned in previous sections. *S. aureus* or *P. aeruginosa* incubated in TSBg sterile media containing only PB was served as control.

**Assessment of Fmoc-F effect on planktonic cells**.

Secondary culture of *S. aureus* or *P. aeruginosa* were assigned the similar treatments as mentioned above. Subsequently, 200 µL of culture in Fmoc-F (1.2 mM) containing media was transferred to 96-well microtiter plate. Different plates were prepared for each time point i.e. 2, 4, 6, 8, 10....till 24 hours. All the plates were properly covered with lids and incubated at 37°C. At the specific time point, planktonic cells were quantified by measuring OD at 600 nm and biofilm cells attached on wells were quantified using crystal violet staining as mentioned previously.

**Anti-biofilm coating test**

Fmoc-F coated sterile coverslips (20 mm diameter) were used as a surface to grow bacterial biofilms. For Fmoc-F coating, coverslips were placed in petri-dish along with 15 mM Fmoc-F solution for overnight incubation. *S. aureus* and *P. aeruginosa* biofilms were grown on the coverslips and attached biomass was quantified by crystal violet assay as mentioned previously.

**Extraction of extracellular matrix (ECM)**

ECM was isolated from bacterial biofilms as described by Chiba *et al*., 2015.[44] Briefly, biofilms grown in conical tubes were centrifuged at 8000 × g for 10 minutes at 4°C. The supernatant was decanted and pellet was washed twice using autoclaved deionized water. The washed pellet was treated with 1.5 M sodium chloride, vortex and kept on rotator for 45 minutes. After that, conical tubes were centrifuged at 5500 × g for 10 minutes at 4°C. The supernatant containing ECM components was transferred to micro-centrifuge tubes, lyophilized using freeze dryer (CHRIST, alpha 2-4 LDplus) and stored for further analysis.

### FTIR analysis of extracted ECM

For studying different components present in biofilm ECM, lyophilized ECM powder was added with sodium-azide which was used as internal standard for spectral analysis. They were mixed properly and put on zinc-selenide crystal of ATR unit of Bruker Tensor 27 IR spectrometer (BrukerOptikGmBH, Ettlingen, Germany). To study the effect of Fmoc-F, ECM grown in presence of 1 mM and 1.2 mM Fmoc-F was evaluated in similar way. For each sample, total 120 scans were recorded in the range of 4000 – 850$cm^{-1}$ with 4$cm^{-1}$ resolution. Spectra were normalized using OPUS software (Opus software, Build 7.2, BrukerOptikGmBH, Ettlingen, Germany).

### Quantification of proteins

Total protein content in extracted ECM was quantified by bicinchoninic acid (BCA) assay using Pierce BCA Protein Assay Kit. Briefly, 25µL of ECM sample was added to 200µL of working reagent as per suggested protocol. The solution was incubated at 37$^oC$ for 30 min until purple colour appeared. 200µL of reaction mixture was transferred to 96 well microtiter plates and OD at 562 nm was recorded. Bovine Serum Albumin (BSA) protein was used to prepare standard curve and protein concentration in ECM samples was quantified using linear curve fitting equation (Figure S2 a).

### Quantification of total polysaccharide

Total polysaccharide concentration in ECM was quantified by using phenol sulphuric acid assay as described by Chiba *et al.* Briefly, 50µL of ECM sample was added to the 125µL concentrated sulphuric acid followed by 25µL of 10% phenol. The tubes were incubated in water-bath (95˚C) for 5 minutes. The reaction mixture was cooled and OD was recorded at 490 nm.[2] Glucose was used to prepare standard curve and carbohydrate concentration was quantified using linear curve fitting equation (Figure S2 b).

**Quantification of DNA**

To quantify the DNA in ECM, a method adapted by Mak *et al.* was used with some modifications. Briefly, 250 µL ECM samples were incubated with DAPI (0.1 X) and fluorescence was recorded at 461 nm.[3] Genomic DNA isolated from *S. aureus* and *P. aeruginosa* was used to prepare standard curve and DNA concentration was quantified using linear curve fitting equation (Figure S2 c, d).

**Thioflavin-T (ThT) binding assay**

ThT binding assay was performed to study the effect of Fmoc-F treatment on amyloid structures in the ECM. Briefly, 195 µL of ECM was taken and incubated with 1.2 mM Fmoc-F (final concentration) for 1 h. Subsequently, 5 µL of ThT (2.5 mM) was added to the mixture and fluorescence intensity was recorded between 400 nm to 650 nm range using spectrofluorometer (LS 55, PerkinElmer). The fluorescence spectra of ThT blank in buffer and ECM blank were also recorded for reference.

### Congo red binding assay

For binding assay, Congo red (CR) solution was prepared in sodium-phosphate buffer (5 mM, pH 7.4) containing sodium chloride (150 mM) and syringe filtered before use. To observe the effect of Fmoc-F, 200 µL of biofilm derived lyophilized ECM (10 mg $mL^{-1}$) was incubated with Fmoc-F (1.2 mM final concentration) for 1 h. The mixture was added with 5 µL of CR solution (7 mg $mL^{-1}$) and kept for 30 min at room temperature. Subsequently, the absorbance spectra of CR was recorded.

### Scanning Electron Microscopy

Biofilms were grown on Fmoc-F (15 mM solution) coated coverslips (10mm) in 24 well plates (described earlier). After 24 hours incubation, media was decanted and biofilm cells were fixed in 2% glutaraldehyde (Merck) for 120 minutes at 4˚C. Samples were then washed thrice with 1X PBS and dehydrated with increasing concentration of alcohol (10 min) and in vacuum desiccator (4 h). All samples were sputter coated with gold and viewed using a scanning electron microscope at 5000X magnification.

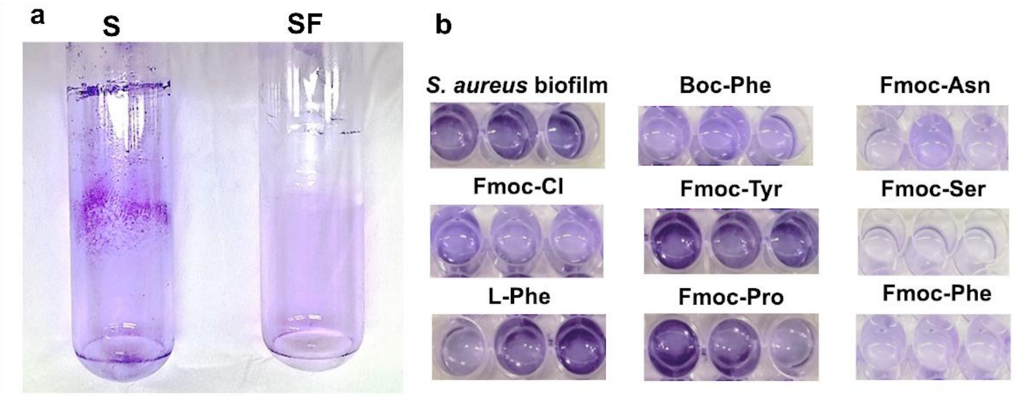


**Figure S1** Anti-biofilm activity of Fmoc- amino acids and L-phenylalanine derivatives against *S. aureus* biofilm as shown by crystal violet staining assay. a) Digital images of biofilm formed by *S. aureus* on glass test-tube walls in the presence or absence of Fmoc-F (1.2 mM). b) Digital images of *S. aureus* biofilm after treatment with Fmoc- amino acids and L-phenylalanine derivatives (1.2 mM). S – *S. aureus* biofilm, SF – *S. aureus* biofilm with Fmoc-F

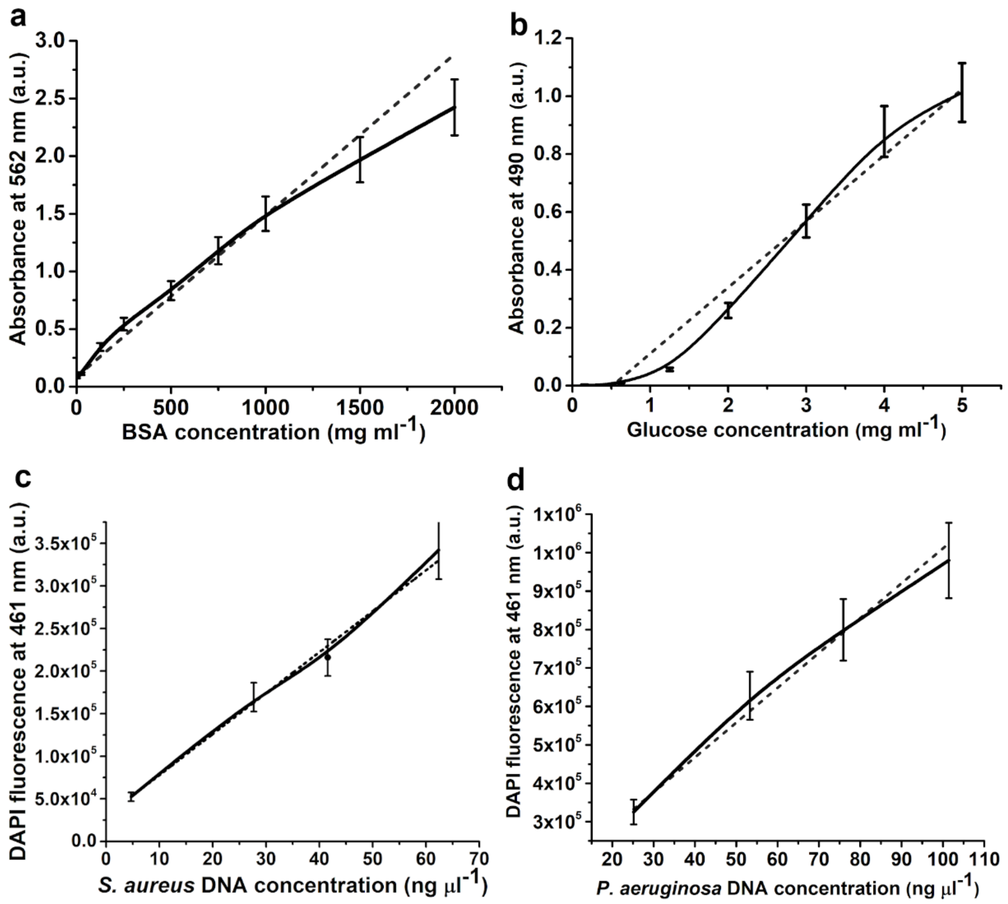


**Figure S2** Standard curve for biochemical assays. a) Bicinchoninic acid assay (BCA) using Bovine serum albumin protein standard where dotted line corresponds to linear fit equation y = 0.0014x + 0.0835, $R^2$= 0.9663 b) Phenol sulphuric acid assay using glucose as carbohydrate standard where dotted line corresponds to linear fit equation y = 0.2283x – 0.117, $R^2$ = 0.966 c) DAPI fluorescence assay using *S. aureus* genomic DNA where dotted line corresponds to linear fit equation y = 4809.3x + 29816.2, $R^2$= 0.993 d) ) DAPI fluorescence assay using *P. aeruginosa* genomic DNA where dotted line corresponds to linear fit equation y = 9064.2x + 104145.6, $R^2$= 0.985

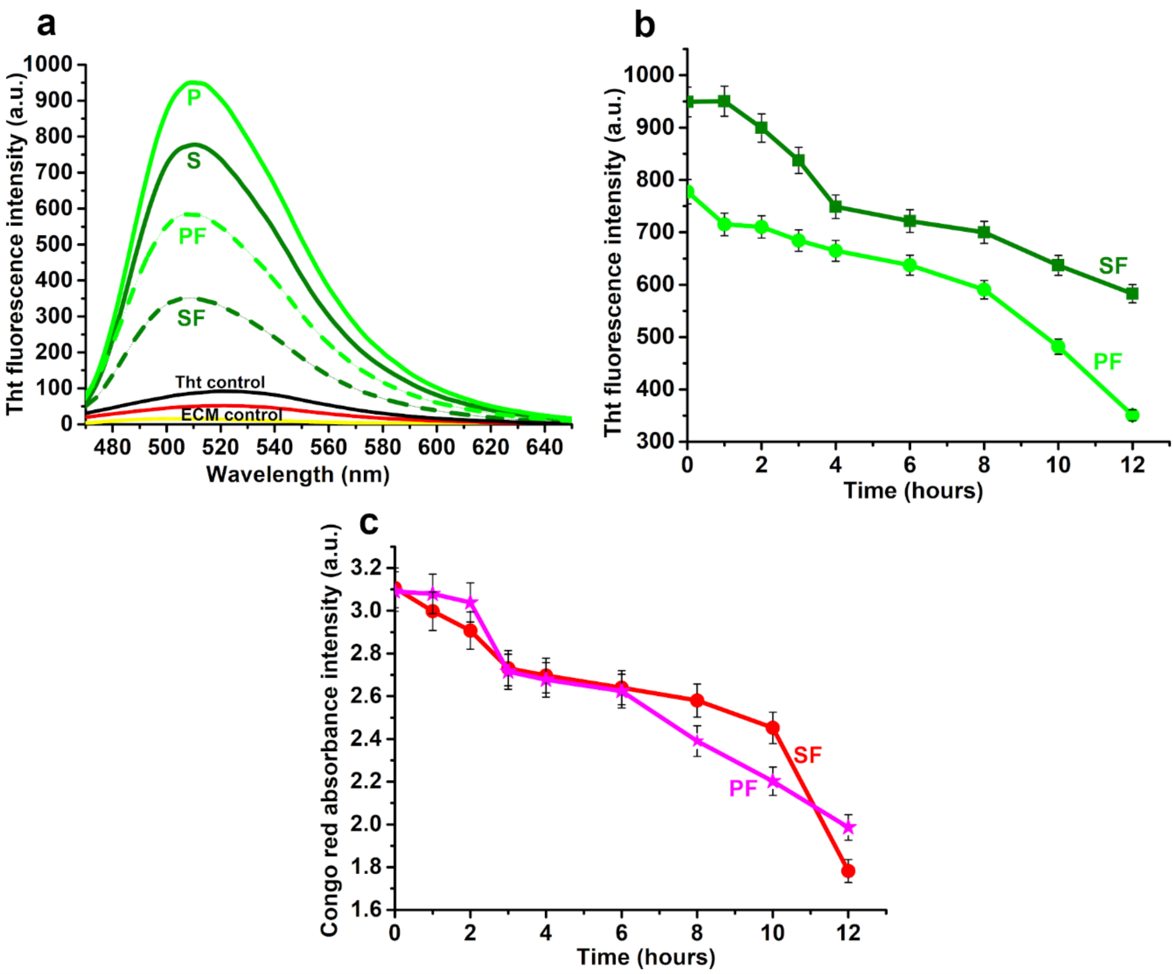


**Figure S3** Effect of Fmoc-F on amyloid structures in ECM. a) Graph showing Thioflavin T fluorescence spectra in presence of ECM extracted from biofilm formed with Fmoc-F (1.2 mM) b) Graph showing decrease in Thioflavin T fluorescence in presence of ECM treated with Fmoc-F (1.2 mM) with increasing time intervals. d) Graph showing reduced Congo red absorbance in presence of ECM treated with Fmoc-F (1.2 mM) with increasing time intervals. S – *S. aureus* ECM, SF – *S. aureus* ECM treated with Fmoc-F, P- *P. aeruginosa*, PF – *P. aeruginosa* ECM treated with Fmoc-F.

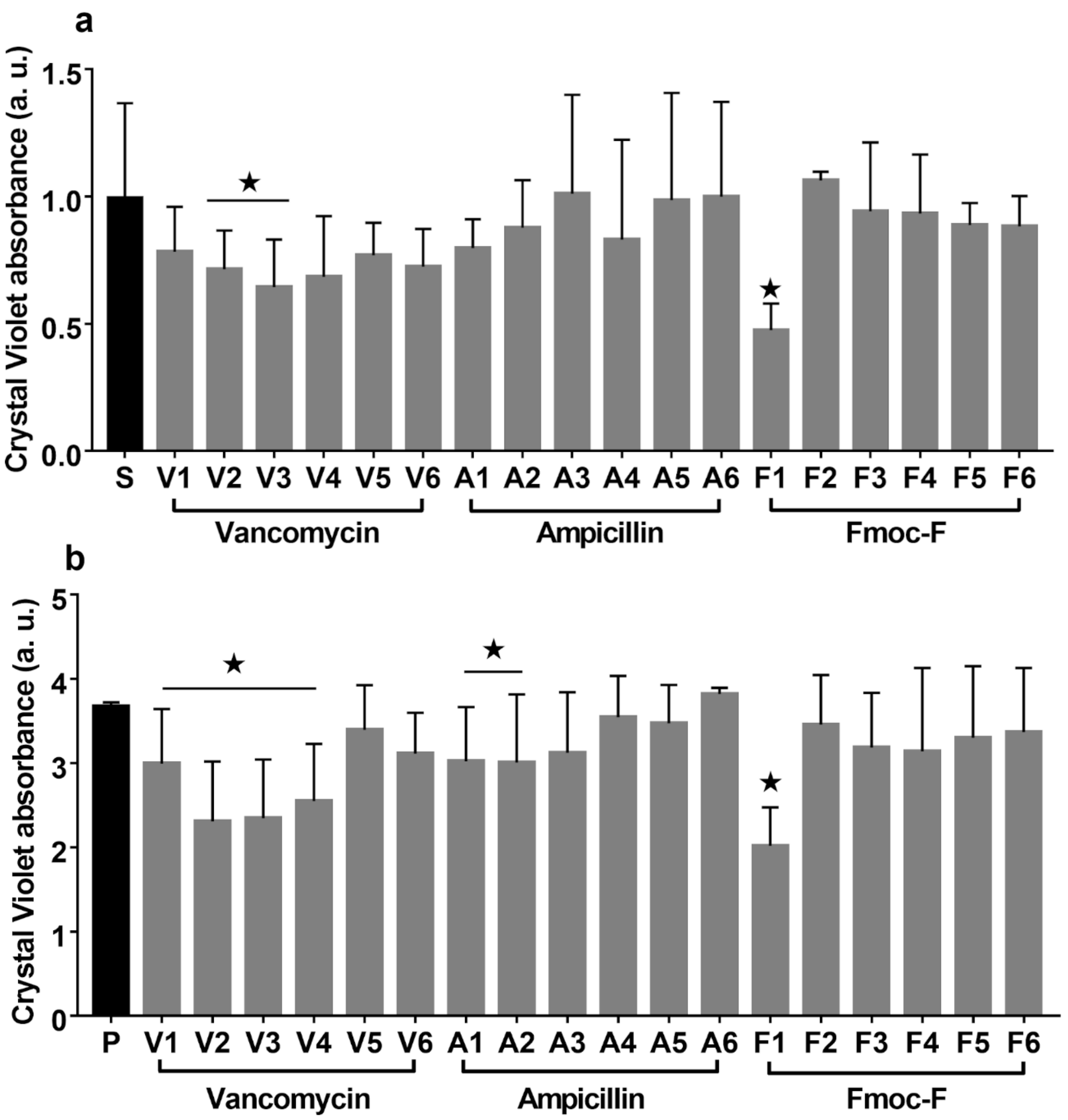


**Figure S4** Eradication of *S. aureus* (a) and *P. aeruginosa* (b) biofilm by treatment with different concentrations of Vancomycin (V1 – 35 µM, V2 – 17.5 µM, V3 – 8.75 µM, V4 – 4.5 µM, V5 – 2.18 µM, V6 – 1.09 µM), Ampicillin (A1 – 143 µM, A2 – 71.5 µM, A3 – 35.7 µM, A4 – 17.8 µM, A5 – 8.9 µM, A6 – 4.45 µM) and Fmoc-F (F1 – 1200 µM, F2 – 600 µM, F3 – 300 µM, F4 – 150 µM, F5 – 75 µM, F6 – 37.5 µM). S – *S. aureus* biofilm, P – *P. aeruginosa* biofilm.
